# Evidence for purifying selection on conserved noncoding elements in the genome of *Drosophila melanogaster*

**DOI:** 10.1101/623744

**Authors:** Berr Tristan, Peticca Aurelie, Haudry Annabelle

## Abstract

Identification of regulatory regions within genomes is a key challenge for understanding the influence of functional traits on species evolution. Development of whole-genome sequencing programs in the early 2000s has been associated with the rise of comparative genomics as a powerful tool for predicting functional regions in genomes, using structural characteristics of DNA sequences. Conserved Noncoding Elements (CNEs, *i.e*. untranslated sequences highly similar across divergent species) were early identified as candidate regulatory regions in metazoans, plants, and fungi. CNEs have been repeatedly discriminated from mutational cold-spots, but only few studies have assessed their functional relevance at a genome-wide scale. In the present study, we used a whole-genome alignment of 27 insect species to build a coherent mapping of CNEs in the genome of *Drosophila melanogaster*. We then exploited polymorphism data from 48 European populations of *D. melanogaster* to estimate levels of nucleotide diversity and adaptive evolution in CNEs. We show here that about 35% of *D. melanogaster* autosomes fall into conserved noncoding regions that exhibit reduced genetic diversity and undergo purifying selection. In addition, we report six insertions of Transposable Elements (TEs) in the genome of *D. melanogaster* showing high levels of conservation across Drosophila species. Half of these insertions are located in untranslated transcribed regions (UTRs) of genes involved in developmental pathways and thus represent potential relics of ancient TE domestication events.

## Introduction

Protein-coding sequences have long been demonstrated to only account for a limited proportion of conserved regions in most genomes (1). 3 to 5% of human DNA has been shown to harbor Conserved Noncoding Elements (CNEs), *i.e*. untranslated sequences highly similar across divergent species (2). In other metazoans, this proportion has been estimated to be much higher (9-15% in *Caenorhabditis elegans* and 25-40% in *Drosophila melanogaster*) (2). Such elements are thought to carry functional roles, even though these functions remain broadly uncharacterized (1). CNEs have indeed been repeatedly distinguished from mutational cold-spots (3), and were shown to exhibit traces of purifying selection in humans (4), *Brassicacea* (5) and *Drosophila* (6). *In vivo* studies have suggested that they often act as enhancers of gene expression (7, 8), but to date no general mechanism accounting for their extreme conservation has been proposed. The informative potential of CNEs regarding regulation – and their apparent influence over human diseases (9, 10) – have made them privileged targets in many comparative genomics approaches, as getting a better grasp on their roles represents a key challenge for understanding both genome evolution and regulatory pathways (1). Genome-wide assessment of conservation and selection thus represents a powerful complement to direct functional studies for characterizing potential regulatory elements in noncoding regions (11).

Identification of CNEs relies on interspecific genomic comparisons (divergence data) (2, 12, 13), while the analysis of their dynamics and evolution is generally performed at an intraspecific scale (polymorphism data) (6, 8). The complementarity of these approaches is at the core of many methods for identifying selection, such as the McDonald & Kreitman test (14) or, more recently, the estimation of deleterious fitness effects (DFE) (15–17). In the fruitfly, *Drosophila melanogaster*, many characterizations of highly conserved elements have been proposed, based on different sets of species and identity criteria (2, 12, 18). However, no systematic study of selection in CNEs combining genome-wide interspecific comparisons with analysis of intra-population polymorphism and selection yet exists in *melanogaster*. Most previously published approaches indeed either focused on small subsets of noncoding regions (6, 19), or neglected to investigate intra-population trends in conserved regions while assessing potential functions of CNEs (2, 12).

Transposable elements (TEs) are mobile DNA elements capable of translocating randomly across an host genome. They have been shown to contribute to the genetic diversity and genome evolution of *D. melanogaster* through interference with regulatory regions (20–22). While most existing studies focus on insertion polymorphism to asses functional roles of TEs (20), comparative genomics offer a way of assessing the effect of past events of TE domestication on genome evolution (23).

In the present study, we used a multiple-species alignment retrieved from the UCSC Genome Browser database, combined with polymorphism data obtained from 48 populations of *D. melanogaster* by the *European Drosophila Population Genomics Consortium* project to (1) identify and characterize CNEs in the genome of *D. melanogaster*, and (**2**) estimate the intensity of purifying selection in CNEs. In addition, we made use of known positions of TE insertions in *melanogaster’*s genome to identify insertions shared across *Drosophila* species, potentially revealing ancient events of TE domestication.

## Material & Methods

### Genomic Datasets

*De novo* search for CNEs in *D. melanogaster* was carried using a whole-genome MULTIZ alignment of 27 insect species and an associated neutral phylogenetic tree, retrieved from the UCSC Genome Browser database (http://hgdownload.soe.ucsc.edu/goldenPath/dm6/multiz27way/). Besides the reference assembly of *D. melanogaster* (BDGP6), the alignment comprised four flies from the *melanogaster* subgroup (*D. simulans, D. sechellia, D. erecta, D. yakuba*), ten flies from the *melanogaster* group (*D. biarmipes, D. suzukii, D. ananassae, D. bipectinata, D. eugracilis, D. elegans, D. kikkawai, D. takahashii, D. rhopaloa, D. ficusphila*), four flies from the *Sophophora* subgenus (*D. pseudoobscura*, *D. persimilis*, *D. miranda*, *D. willistoni*), four flies from the *Drosophila* genus (*D. virilis*, *D. mojavensis*, *D. albomicans*, *D. grimshawi*), two *Diptera* (*Musca. domestica, Anopheles. gambiae*), one *Hymenoptera* (*Apis. mellifera*) and one *Coleoptera (Tribolium. castaneum*). Divergence between *Drosophila* species was estimated to range from 25 to 40 My (24) (vs >300 My for the whole alignment (25)).

Polymorphism data in 48 European populations of *D. melanogaster* was obtained from the *European Drosophila Population Genomics Consortium* (*DrosEU*). Samples consisted in: nineteen populations from Ukraine, four populations from Germany and the United Kingdom, three populations from Austria, Finland, and France, two populations from Denmark, Spain, Switzerland, and Turkey and one population from Cyprus, Portugal, Russia, and Sweden. Paired-end Illumina Pool-seq analysis was performed on thirty-three to forty male diploid genomes for each sample. The output reads were subsequently trimmed, aligned on the reference sequence of *melanogaster*, and filtered in order to retain robust polymorphic sites (Kapun et al. 2018).

### Identification of conserved elements

We made use of the R-friendly version of PHAST (PHylogenetic Analysis with Space-Time models, release 1.6.5 (26)) to retrieve discrete conserved elements in *D. melanogaster*’s genome. PHAST takes a multiple-species alignment combined with a phylogenetic tree for building a phylogenetic-Hidden Markov Model of sequence evolution (PhyloFit). This model is then used to assign basewise conservation scores along a region of interest. Finally, discrete conserved blocks can be identified, based on expected values of coverage and length for conserved elements (PhastCons).

The software was run along the 2L, 2R, 3L, 3R, and X chromosomes of *D. melanogaster* (due to poor sequence conservation outside of the *melanogaster* subgenus, chromosomes 4 and Y were not searched for CNEs). Because indels and gaps in an alignment are treated as missing data by PHAST, we investigated the effect of phylogenetic distance on the quality of PhastCons outputs (the more distant the species, the more regions are mistaken for conserved elements because of mis-alignment). The final set of predicted conserved elements was obtained by running PhastCons on a subset of fifteen species from the *melanogaster* group, with an expected coverage of 0.3 and an expected length of 45 bp.

### CNE filtering

Based on *D. melanogaster*’s genome annotations (Ensembl 87, Refseq genes), predicted conserved elements overlapping with coding regions were discarded. The remaining CNEs were filtered against blocks shorter than 6 bp or that were not shared by at least thirteen of the fifteen species (due to *D. ananassae* and *D. bipectinata* having rapidly diverged from the rest of the *melanogaster* group). To_investigate for potential roles of CNEs, their positions were mapped against the annotated genome of *D. melanogaster*: CNEs that fell into several types of annotations were split into smaller blocks with a single identity. The classes of CNEs that were retained comprised: CNEs in noncoding RNAs, in proximal regions of genes (<500 bp away from the borders of a transcribed region), in 5’ UTRs of genes, in 3’ UTRs of genes, in introns, and in distal intergenic regions (>500 bp away from the borders of transcribed regions). Using this newly built annotation set of CNEs, we tested for enrichment of annotated regions in conserved elements (bilateral two-samples proportion test) by comparing the composition of CNEs in terms of annotation types (UTRs, Introns, ncRNAs, …) with that of the whole genome of *D. melanogaster*. Distribution of CNEs along *melanogaster*’s chromosomes was investigated for identifying potential clusters of conserved regions. CNEs were also tested (chi-squared comparison of multiple distributions) for differences in size and GC-content between different annotation types.

### Polymorphism estimation in CNEs

Using a cartography of biallelic SNPs in the 48 DrosEU populations (chromosomes 2L, 2R, 3L, 3R, and X), we measured genetic diversity in each sample by computing polymorphism estimators: Tajima’s *π*, Watterson’s *θ* and Tajima’s *D* (27–29). Due to SNPs having been filtered for keeping robust polymorphic positions, corrected versions of polymorphism estimators for naïve Pool-seq data (27) were not used. we compared levels of polymorphism within CNEs with those in fourfold- (4D) and zerofold-degenerate (0D) sites, respectively used as a proxy fo neutral and highly constrained evolution. The estimators were first computed on each chromosome separately and then averaged in order to get a single value per category (CNEs, 4D, 0D) and sample. Correlation between estimates of *π, θ* and *D* and geographic characteristics of the samples (latitude, longitude) were then tested. Due to significant variation in sequencing quality, the recovery of low-frequency variants fluctuated widely across populations. This strongly affected the estimation of *θ* (and, subsequently, of Tajima’s *D*) and hindered the assessment of population-specific evolutionary trends.

### Construction of Allele Frequency Spectra

Derived allele frequency spectra (DAFS) were built for polymorphic sites (CNEs, 4D, 0D) for each population using two outgroups (*D. sechellia, D. erecta*) to assess the ancestral/derived state of an allele (*e.g*., if a SNP harbored an A/T polymorphism, A or T was considered ancestral only if present in the two outgroups). Sites with ambiguous allele state were discarded. Populations were then investigated individually for assessing the distribution of allele frequencies. A DAFS comprising all populations was eventually built by summing the 48 individual frequency spectra in order to estimate general selective constraint over CNEs. Computing population-wise DAFS allowed to correctly estimate the frequencies of alleles whose prevalence fluctuate across samples (an allele may be found at a low frequency in a population while being strongly represented in other samples).

### Measuring the proportion of adaptive substitutions in CNEs

Keightley & Eyre-Walker (15) have developed the software DFE-alpha for measuring the proportion of adaptive substitutions (α) in a putatively selected region using allele frequency spectra. We made use of two of their programs (release 2.15): est_dfe for measuring the strength of deleterious fitness effects and est_alpha_omega for estimating the value of α in CNEs. Runs of est_dfe and est_alpha_omega were performed on the different classes of CNEs using the combined DAFS for all populations, and assuming a constant population size for *melanogaster* samples. In addition to allele frequency spectra, estimation of alpha requires counts of divergent sites in selected regions, which were recovered using *D. sechellia* and *D. erecta* as outgroups (polymorphic sites contributing to the allele frequency spectra were not considered when estimating divergence).

### Search for ancient transposition events

We used Flybase (release 6.15) to retrieve known insertions of Transposable Elements along the genome of *D. melanogaster* (5392 coordinates). Overlaps with previously identified CNEs were recovered and filtered against fragments shorter than 6 bp. The remaining elements were classified with respect to their transposon families, length, and level of conservation. Neighboring regions of insertions conserved over more than 100 bp were subsequently investigated for assessing potential functions, by examining nearby genes (<500 bp away from the borders of an insertion).

## Results

### Effects of phylogenetic distance on PhastCons detection power

Assessing the effect of phylogenetic distance on PhastCons detection power revealed that species past a divergence threshold (for this case, outside of the *Drosophila* genus) did not improve the detection of conserved elements, due to poor genomic resemblance with the target species. (*D. melanogaster*) When only considering blocks shared by a minimum number of species (in order to remove regions containing indels and gaps), we found PhastCons performances to be nearly identical when the program was run on all species (27 insects) or on a subset of 15 species from the *melanogaster* group. However, recovery of conserved regions dramatically decreased below a certain divergence threshold (PhastCons run within the *melanogaster* subgroup, 5 species). This is likely due to an inaccurate discrimination of conserved/non-conserved regions when little divergence has affected the genome sequence, despite the program’s effort for taking phylogeny into account when identifying conserved elements.

### Identification of CNEs in D. melanogaster

PhastCons runs on *D. melanogaster*’s chromosomes 2L, 2R, 3L, 3R, and X, and subsequent filtering (see *Material & Methods*) yielded approximately 1,115,000 conserved noncoding regions encompassing 41.6 Mb (Table 1). Most CNEs were found in introns (47% of total coverage) and distal regions more than 500 bp away from a gene (28%), with untranslated transcribed regions (UTRs), noncoding RNAs (ncRNAs), proximal regions and pseudogenes sharing the remaining 25 %. The median size of CNEs (all annotation types pooled) was 23 bp, with a clear majority (93%) being shorter than 100 bp. Despite apparent homogeneity in size distribution and GC-content across annotation types, all classes of CNEs were found to have significantly different lengths and GC% (chi-squared comparison of multiple distributions, P-values < 0.001). This may be a consequence of large sample size dramatically increasing the power of statistical measurements (see *Discussion*).

**Table 1.** Summary of the filtered CNE dataset

| *Type* | *No. of CNEs* | *Total size (bp)* | *Median size<sup>1</sup> (bp)* | *MAD<sub>size</sub><sup>2</sup>* | *Sk<sub>size</sub><sup>3</sup>* | *GC%* | *MAD<sub>GC%</sub><sup>2</sup>* | *Sk<sub>GC%</sub><sup>3</sup>* |
| --- | --- | --- | --- | --- | --- | --- | --- | --- |
| *All* | *1,115,754* | *41,583,499* | *23* | *19.3* | *4.49* | *39.4* | *13.0* | *-0.200* |
| *ncRNAs* | *48,221* | *1,816,917* | *24* | *20.8* | *3.39* | *41.6* | *12.5* | *-0.268* |
| *Proximal* | *106,425* | *3,282,013* | *19* | *14.8* | *3.34* | *40.0* | *14.8* | *-0.190* |
| *5' UTRs* | *61,788* | *1,879,665* | *19* | *14.8* | *6.05* | *40.7* | *13.7* | *-0.211* |
| *Introns* | *527,942* | *19,594,612* | *23* | *19.3* | *3.56* | *39.1* | *12.4* | *-0.221* |
| *3' UTRs* | *94,020* | *3,441,561* | *21* | *17.8* | *8.14* | *32.0* | *13.1* | *0.179* |
| *Pseudogenes* | *478* | *16,297* | *18* | *14.8* | *2.10* | *44.9* | *13.4* | *-0.542* |
| *Distal* | *276,880* | *11,552,434* | *26* | *23.7* | *3.46* | *41.2* | *11.6* | *-0.284* |

Types: noncoding RNAs, intergenic regions within 500 bp of a gene (proximal), untranslated transcribed regions, introns, pseudogenes, intergenic regions more than 500 bp away from a gene (distal).
<sup>1</sup>Median values were deemed more representative than mean estimates due to the asymmetric size distribution of conserved blocks and the presence of rare long (>1,500 bp) CNEs.
<sup>2</sup>For similar reasons, MAD (*Median Absolute Deviation*) was used instead of standard deviation (strongly affected by extreme values) or 95 % confidence intervals (due to large sample size).
<sup>3</sup>Adjusted Fisher-Pearson coefficient of skewness: a positive value indicates that the sample distribution is skewed right, *i.e.* that the right tail is long relative to the left tail (opposite for negative value).

CNEs located within pseudogenes were not considered in further analyses, due to their rarity (<1‰ of total coverage).

### Enrichment of functional regions in CNEs

Comparing the composition of CNEs with that of the whole genome of *D. melanogaster* (two-samples proportion test) revealed UTRs and ncRNAs to be significantly enriched in conserved regions (Fig.1, P-values < 0.001). Compared to random distribution of conserved positions, sites in UTRs were more often conserved (+27% for 3’ and +18% for 5’ UTRs). Introns and intergenic regions, on the opposite, appeared depleted in conserved positions, with introns, proximal, and distal regions containing respectively −13%, −23% and −24% less conserved bases than expected under a random distribution. Finally, overall distribution of CNEs along chromosomes revealed no clusterization of conserved regions.

**Figure 1:**
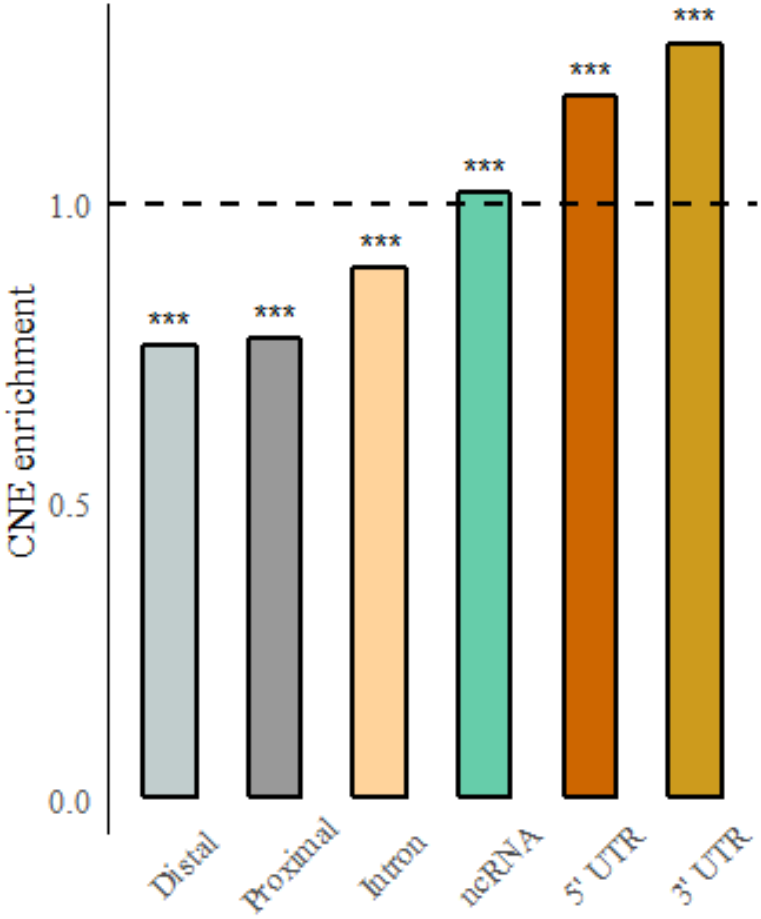
Relative enrichment in CNEs by annotation type. ***: P-value < 0.001 (Two-samples proportion tests between CNEs composition and genome composition).

### CNEs conservation across species

CNEs conservation within the different *Drosophila* clades was found to reflect phylogenetic distance (Fig.2). CNEs exhibited strong conservation (>90% of CNEs with >80% sequence identity) within the *melanogaster* subgroup (5 species closest to *D. melanogaster*). Overall conservation appeared reduced within the *melanogaster* group, especially due to species having rapidly diverged from *D. melanogaster* (*D. bipectinata, D. ananassae, D. kikkawai*). Only 25% of CNEs appeared shared by *Sophophora* flies outside of the *melanogaster* group, and nearly no CNEs (<1‰) were found shared outside of the *Drosophila* genus.

**Figure 2:**
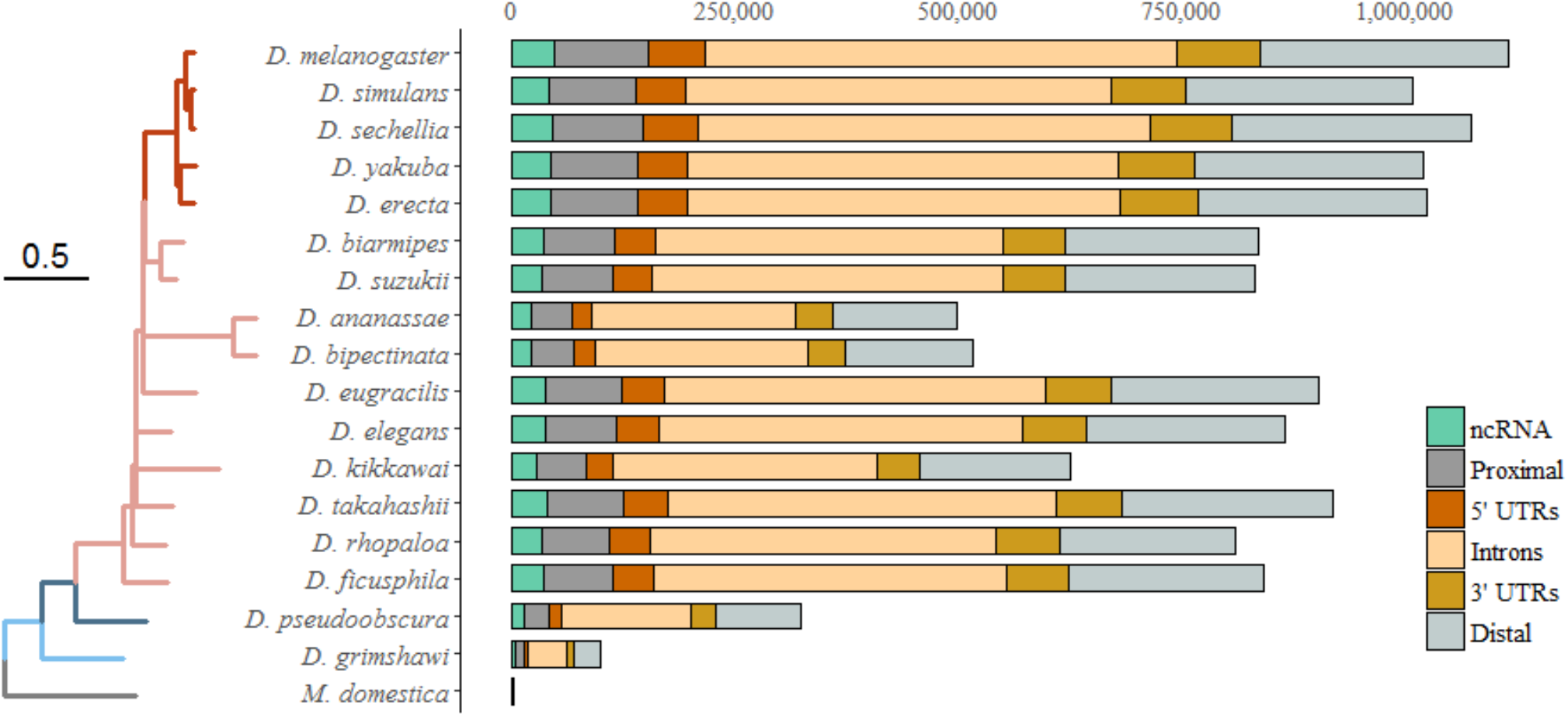
Conservation of CNEs across drosophilid species. Counts of CNEs shared with *D. melanogaster* (>0.8 sequence identity) are presented by species, with respect to phylogenetic distance. Sampled species belong to the melanogaster group (15 species), either in (red, 5 species) or outside of (pink, 10 species) the melanogaster subgroup. Counts for *D. pseudoobscura* (Sophophora subgenus, darkblue), *D. grimshawi* (Drosophila genus, lightblue) and *Musca domestica* (Muscidae) indicate conservation outside of the *melanogaster* group. Tree scale: Number of substitutions per site.

### Reduced nucleotide diversity in CNEs

In all DrosEU populations, CNEs exhibited a level of nucleotide diversity (Tajima’s π) intermediate between 0D and 4D sites (Fig.3.A). 4D sites (neutral expectation) showed the highest values (1.03 × 10^-2^ ± 0.02 across all samples pooled) and 0D sites (constrained expectation) the lowest (2.04 × 10^-3^ ± 0.27). Overall, nucleotide diversity within CNEs appeared closer to that in 0D sites (3.27 × 10^-3^ ± 0.14). No significant correlation of *π* with latitude or longitude could be uncovered.

**Figure 3:**
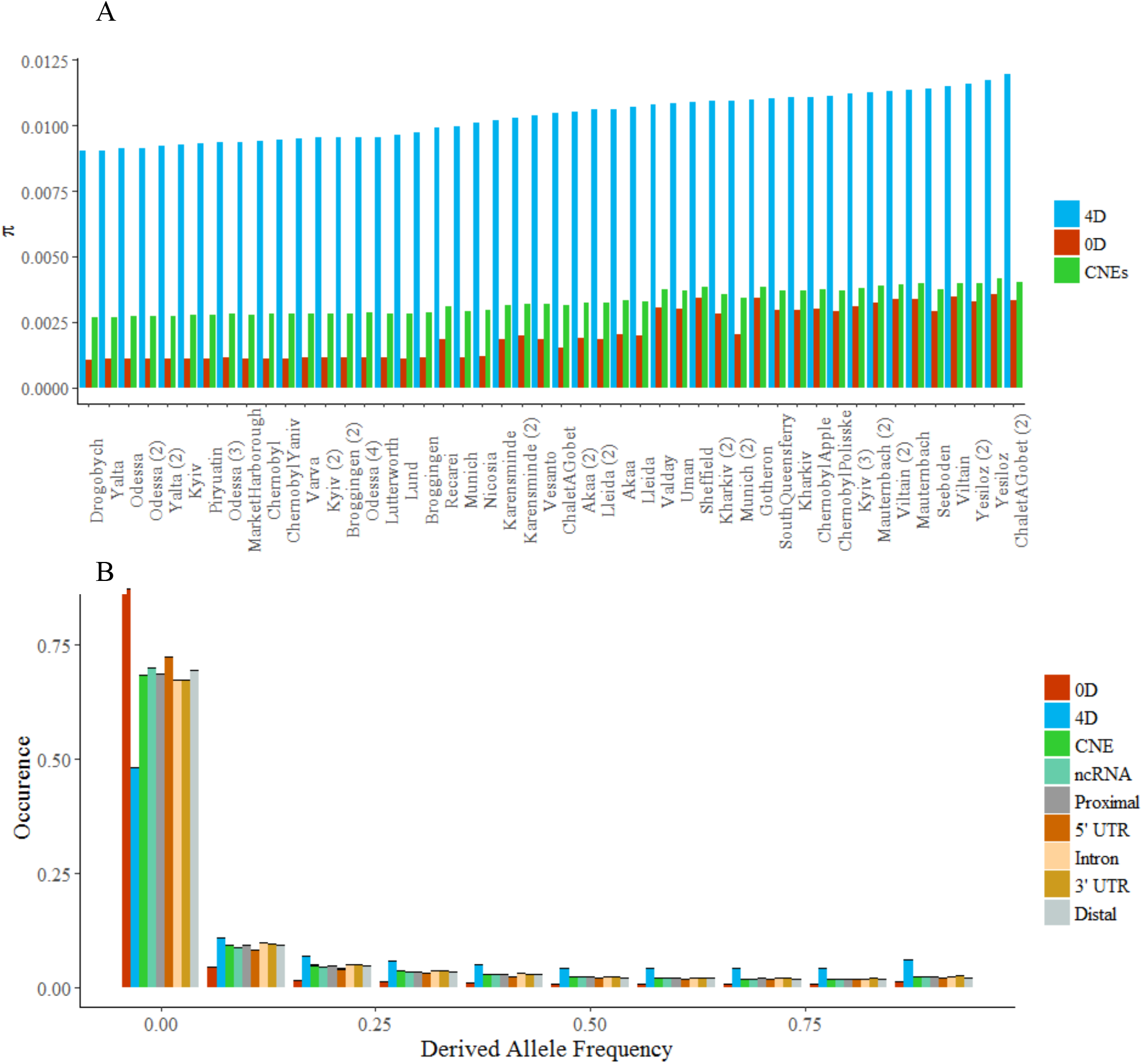
Evidence for reduced polymorphism and purifying selection in CNEs. A: Distribution of Tajima’s π across 48 European populations of D. melanogaster in CNEs (green), zero-fold (0D, red) and fourfold (4D, blue) degenerate sites. Estimates of π were computed from a set of ~5,360,000 biallelic SNPs (polymorphic sites) and a mapping of robust monomorphic positions. B: Derived Allele individual DAFS) in different classes of CNEs, 0D ans 4D sites. Black horizontal lines indicate bootstrapped confidence intervals (95% - 100 replicates).

### Purifying selection in CNEs

DAFS within polymorphic positions of *D. melanogaster’*s genome revealed a significant excess of rare variants (frequency < 0.1) in both 0D sites and CNEs, compared to 4D sites (Fig.3.B). This excess indicates that mutations in CNEs and 0D sites tend to increase less in frequency than those in 4D sites, *i.e*. that they undergo purifying selection. 0D sites displayed the highest level of selective pressure, while CNEs showed intermediate intensity of negative selection with little variation across classes.

### Adaptive substitutions in CNEs

Estimates of α, the fraction of nucleotide divergence driven by positive selection, were computed in the different classes of CNEs using the software DFE-alpha (*see Material & Methods*) on newly-built DAFS. When all classes of CNEs were pooled, α was estimated at 0.29, *i.e*. 29% of nucleotide divergence in CNEs is expected to be driven by positive selection. α was found highest in UTRs (α_5’ UTRs_ = 0.49, α_3’ UTRs_ = 0.40) and ncRNAs (α_ncRNAs_ = 0.42), intermediate in proximal regions (α_proximal_ = 0.33), and lowest in distal regions and introns (α_distal_ = 0.28, α_introns_ = 0.24).

### Ancient transposition events

Matching the positions of the previously identified CNEs against known TEs insertions in the reference genome of *D. melanogaster* yielded 284 overlapping regions encompassing 8510 bases (median size: 18 bp). Six insertions were found conserved over more than 100 bp (in CNEs only), among which five were partially located in UTRs of genes (Table 2). These six insertions were present in the fifteen species of the *melanogaster* group used for identifying conserved elements.

**Table 2.** Characteristics of TE insertions conserved over more than 100 bp

| *TE (insertion<sup>\*</sup>)* | *Size (bp)* | *No. of bases in CNEs* | *Annotation* | *Neighbor genes* |
| --- | --- | --- | --- | --- |
| **2R** |  |  |  |  |
| *roo* (850) | 8921 | 4575 | 5' UTR, intron, proximal | *grh*<sup>1</sup> |
| *INE-1* (3460) | 1751 | 223 | 5' UTR, proximal | *predicted gene*<sup>2</sup> |
| *INE-1* (1756) | 1675 | 188 | 3' UTR | *pld*<sup>3</sup> |
| *HMS-Beagle* (3102) | 415 | 333 | 5' UTR | *predicted gene*<sup>2</sup> |
| **3L** |  |  |  |  |
| *FB* (1604) | 8159 | 738 | 5' UTR, intron, 3' UTR | *spock*<sup>4</sup>, *predicted gene*<sup>2</sup> |
| **3R** |  |  |  |  |
| *HMS-Beagle* (4853) | 195 | 192 | 5' UTR | *ssdp*<sup>5</sup> |
| **X** |  |  |  |  |
| *roo* (53) | 1194 | 166 | *distal* | - |

<sup>\*</sup>Insertion numbers as referenced in Flybase (release 6.15).
<sup>1</sup>*grh* (grainyhead): transcription factor (interacts with *ddc*, *engrailed*, *ultrabithorax*).
<sup>2</sup>Predicted genes referenced in Flybase, either identified through genome-wide screening for active coding regions or sequences that align with mRNAs recovered by RNAseq.
<sup>2</sup>*spock*: calcium-transporting ATPase.
<sup>3</sup>*ssdp* (single-stranded dna-binding protein): ARN polymerase II promoter (predicted).
<sup>4</sup>*pld* (phospholipase D): phospholipase.
Source: Flybase, UniProtKB.

All statistical analyses were performed using R 3.3.1.

## Discussion

### Limits of statistical significance in very large datasets

Genome-wide datasets (alignments, annotation files, polymorphism data…) are generally characterized by a very large number of entries, leading most statistical measurements to exhibit extreme significance (30). This phenomenon is linked to large samples drastically increasing the power of statistical tests, even if the practical significance of the considered effects (*i.e*. the interpretation that can be drawn from the observed trend) is nearly nonexistent. Thus, many of the tests performed on CNEs, such as the estimations of differences in size and GC% between annotation types, produced extremely low p-values, even though the observed differences seemed far too weak to be interpreted as consequences of a biological effect. Both standard errors and bootstraps are strongly affected by this “large-sample size issue” (30).

### Influence of phylogenetic distance on PhastCons detection power

Since its original publication (2), the PhastCons program (now implemented in the PHAST software (26)) has been widely used for retrieving discrete conserved elements in many metazoans and plants. Most species identified in the UCSC Genome Browser database have an associated PhastCons track of predicted conserved regions. Because of the questionable handling of indels and gaps by PhastCons (read as missing data), we chose not to use the existing set of predicted conserved elements available in the UCSC database for *D. melanogaster* (http://hgdownload.soe.ucsc.edu/goldenPath/dm6/phastCons27way/), identified using a multiple-species alignment of 27 insects. When examining sequence conservation within elements from the UCSC dataset, we determined that the most distant species have very little influence on the identification of conserved blocks. Moreover, large phylogenetic distance with *D. melanogaster* increases the risk of mis-alignment of non-homologous regions being interpreted as traces of conservation, resulting in some predicted conserved elements being only shared by with the most distant species.

### CNE content in Drosophila

The number and coverage of predicted CNEs in chromosomes 2L, 2R, 3L, 3R, and X of *D. melanogaster* identified in this study is fairly consistent with existing measurements of conservation in *Drosophila* (1, 2, 19). In the original PhastCons paper (2), Siepel & al. estimated that 27.8 to 39.8 % of *melanogaster*’s genome fell into conserved noncoding regions, using a 4-species alignment of insects (*D. melanogaster, D. yakuba, D. pseudoobscura, A. gambiae*). We evaluated here that about 35% of bases within the investigated chromosomes were comprised in CNEs. Due to numerous revisions of gene annotation in *melanogaster*’s genome since the original Flybase report (31), the proportion of conserved elements falling into unannotated regions has been repeatedly revised downward (25% in this study vs ~40% in Siepel’s paper), mostly to the benefit of novel-discovered genes (32). Contrary to CNEs in humans (33), conserved noncoding regions in *Drosophila* were found to be evenly distributed across the genome, with no trace of local clusterization. This is likely linked to the greater compaction of homogeneity of *D. melanogaster*’s genome compared to human (34).

The observed enrichment of noncoding transcribed regions (UTRs, ncRNAs) in CNEs, opposed to the reduced conservation of introns and intergenic regions is coherent with known distribution of CNEs in *Drosophila* (2, 19), human (4), and *Arabidopsis* (5). At the genome-wide scale, it supports the hypothesis that conservation in CNEs is partially driven by functional importance (1, 3, 6): UTRs are known to be involved in gene expression and regulation and noncoding RNAs are among the most documented examples of regulatory elements in genomes (35)

### Polymorphism estimation European populations

The European consortium DrosEU recently generated Pool-Seq data from wild-caught flies from diverse sampling sites (Kapun et al. 2018). We used SNP variants detected in 48 samples from more than 30 localities all across Europe for our polymorphism analyses. Recovery of low-frequency SNP variants in the DrosEU populations was strongly affected by sequencing bias. Read coverage among samples fluctuated widely depending on the sequencing run (5 in total), resulting in an uneven recovery of rare variants across populations, as the quality thresholds used by the DrosEU Consortium discarded low-frequency alleles more easily when sequencing depth was small. While nucleotide diversity (Tajima’s *π*) is only marginally affected by rare variants, Watterson’s estimator of genetic diversity (*θ*) is heavily influenced by their presence/absence. This difference in handling rare variants is used when computing Tajima’s *D* to estimate excess/rarity of low-frequency variants, allowing to infer recent demographic events and/or patterns of selection. Here, the uneven recovery of rare alleles across populations rendered impossible a correct estimation of Watterson’s θ, and subsequently of Tajima’s *D*. Assessing differences in nucleotide diversity (and later in allele frequency spectra) between classes of SNPs (CNEs, 4D, and 0D sites) could however still be performed, due to low-frequency variants being evenly omitted in the genome.

Nucleotide diversity in CNEs was found close to that in 0D sites, and much inferior to estimates of *π* within 4D sites. This is coherent with CNEs exhibiting high sequence similarity across divergent species (seemingly conserved regions are expected to be less polymorphic than the rest of the genome) and with existing estimations of polymorphism in noncoding regions (6) of *melanogaster*. This result alone, however, cannot be interpreted as a consequence of purifying selection: conservation and reduced polymorphism in CNEs may simply owe to lower mutation rates.

### Purifying selection in CNEs

Purifying selection in genomes can be discriminated from reduced mutation rates by examining the distribution of derived allele frequencies in a supposedly constrained region, compared to a neutral expectation (3, 36). Negatively selected sites tend to exhibit an excess of rare variants compared to neutrally evolving positions, due to new mutations in constrained regions being unlikely to spread. In this study, we estimated that CNEs in the genome of *D. melanogaster* are less constrained than 0D sites, but still exhibit an excess of rare variants compared to 4D positions. These results confirm that conservation in CNEs is driven by selective constraint and not by local variation in mutation rates (3, 6, 19, 37), and provide a strong validation of their functional importance in genomes. Selection intensity appeared highly similar across the different classes of CNEs, likely illustrating the general homogeneity of conserved elements predicted by PhastCons.

To seek further confirmation of the functional roles of CNEs, we estimated the proportion of substitutions between *D. melanogaster and* two closely related species (*D. sechellia, D. erecta*) driven by positive selection. In agreement with the work of Andolfatto (6), we found UTRs and ncRNAs to be the noncoding regions most subject to adaptive selection, with more than 40% of divergence being positively selected.

### Ancient transposition events

Transposable elements are known to contribute to genetic diversity in populations of *D. melanogaster*, and have been repeatedly shown to influence genome evolution (20, 38). Studies on TE polymorphism have demonstrated that insertions within regulatory regions of genes can present adaptive potential (21, 22) and become fixed through positive selection. Little is known, however, on past domestication events that may have affected gene expression in *melanogaster*’s genome, as most functional characterizations of TEs have been conducted using polymorphism of TE insertions (*i.e*. an already fixed insertion will not appear to affect gene regulation). In mammals, 16% of eutherian-specific CNEs have been estimated to originate from ancient transposition events (23), but to my knowledge no similar assessment of TE contribution to CNEs has been proposed in *Drosophila*. In the present study, we reported six TE insertions conserved over more than 100 bp in noncoding regions, all of which were shared by the 15 species considered in the *melanogaster* group. Three of these insertions are partially located in UTRs of genes involved in developmental processes (*grh, pld, ssdp*). There is therefore strong support for proposing that these “conserved noncoding insertions” became domesticated in the common ancestor of the *melanogaster* group (15-20 My (24)).

## Author contributions

CNEs identification and characterization, as well as polymorphism analyses for identifying selection were conducted by Tristan Berr, under the guidance of Annabelle Haudry, Phd. Genomic datasets were provided by the UCSC Genome Browser database and the European Drosophila Population Genomics Consortium.

## Acknowledgements

The author thanks M. Hubisz for help in exploring the subtleties of the *rphast* package for R, M. Kapun for advice and information on *DrosEU* polymorphism data and C. Vieira-Heddi for helpful discussions.

*Rphast* is a R package developed by M. Hubisz & al. for analyses in comparative and evolutionary genomics.

*DFE-alpha* is a software developed by A. Eyre-Walker and P. Keightley for estimating deleterious fitness effects and characterizing adaptive substitutions using site frequency spectra.

